# Pathological variants in *HPDL* cause collapse of the neuro-glial unit during human cortical maturation

**DOI:** 10.64898/2026.06.09.731096

**Authors:** Matteo Baggiani, Michela Giacich, Valentina Naef, Filippo Maria Santorelli, Devid Damiani

## Abstract

Biallelic variants in *HPDL* cause a severe neurological disorder in childhood, but the mechanisms linking early developmental abnormalities to later cortical degeneration remain unclear.

Here, using long-term iPSC-derived human cortical cultures derived from four patients, we investigated the late consequences of HPDL deficiency through quantitative immunofluorescence and bulk RNA-seq.

We found that HPDL-deficient cortical cultures undergo progressive synaptic impairment, with marked loss of PSD95-positive postsynaptic puncta despite largely preserved general neuronal maturation markers. This phenotype is accompanied by a profound reduction in astrocytes and oligodendrocytes, together with increased neuronal apoptosis and transcriptional dysregulation of genes linked to extracellular matrix organization, cellular stress, and neurodegeneration.

Unexpectedly, late-stage mutant cultures also show reactivation of early developmental programs, including aberrant expression of NEUROD4 and other proneural regulators, suggesting instability of cell identity during cortical maturation. Together, these findings support a model in which HPDL deficiency first perturbs cortical developmental timing and later drives collapse of the neuro-glial unit, linking premature neurogenesis to synaptic failure, glial loss, and progressive neurodegeneration.

## Introduction

Biallelic pathogenic variants in *HPDL* (4-hydroxyphenylpyruvate dioxygenase-like) cause a rare neurodegenerative disease ranging from neonatal encephalopathy with intractable epilepsy and severe developmental delay to childhood- or adolescent-onset hereditary spastic paraplegia with progressive lower-limb spasticity and white-matter abnormalities (Brunetti et al., 2021; Nonaka-Kinoshita et al., 2013). Clinically, patients frequently present early hypotonia, developmental arrest, movement disorders, seizures, corticospinal tract problems, and neuroimaging findings such as white matter abnormalities, and cortical atrophy (Alecu et al., 2025).

At the molecular level, HPDL is a mitochondrial protein that localizes to the intermembrane space and belongs to the vicinal oxygen chelate superfamily, containing two HPPD-like domains with a conserved iron-binding motif (Alecu et al., 2025; Husain et al., 2020). Although initially of unknown function, recent studies revealed that HPDL catalyzes the conversion of 4-hydroxyphenylpyruvate into 4-hydroxymandelate (4-HMA), precursors of 4-hydroxybenzoate (4-HB) in Coenzyme Q₁₀ (CoQ₁₀) alternative synthesis pathway (Banh et al., 2021). In *Hpdl*-null mice, loss of HPDL leads to primary CoQ deficiency in the brain, impaired electron flow through complexes I and II, cerebellar mitochondrial fragmentation, and lethal encephalopathy with seizures and spasticity by postnatal day (Ghosh et al., 2021; Wiessner et al., 2021). Remarkably, oral supplementation with 4-HMA or 4-HB partially restores brain CoQ, rescues electron transport, normalizes Purkinje-cell function and cerebellar histology, and allows almost all *Hpdl*^-/-^ mice to reach adulthood. Moreover, in a child with pathological *HPDL* variants associated with spastic ataxia features, 4-HB treatment similarly stabilized neurological symptoms over several months (Shi et al., 2025). These data position HPDL as a key regulator of brain CoQ homeostasis and mitochondrial respiratory-chain function during development (Chou et al., 2018; Fame and Lehtinen, 2021).

Another important finding is the recognition of mitochondrial metabolism and redox signaling as central regulators of neural stem cell (NSC) maintenance, neurogenesis, and cortical development (Brunetti et al., 2021; Garone et al., 2024; Iwata and Vanderhaeghen, 2021; Iwata et al., 2023; Lorenz and Prigione, 2017; Pilaz et al., 2016). NSCs in embryonic and adult brain undergo a metabolic shift from predominantly glycolytic flux to oxidative phosphorylation (OxPhos) as they commit to a neuronal fate, accompanied by changes in mitochondrial morphology (from fused to fragmented), membrane potential, and reactive oxygen species (ROS) production (Armat et al., 2025; Iwata and Vanderhaeghen, 2021; Iwata et al., 2023). Mitochondrial dynamics and ROS act as instructive signals: transient mitochondrial fragmentation and moderate ROS levels promote neuronal differentiation via NRF2- and Wnt/β-catenin-dependent pathways, whereas sustained mitochondrial dysfunction or excessive ROS deplete the NSC pool, impair adult neurogenesis, and ultimately cause cognitive decline (Iwata and Vanderhaeghen, 2021; Khacho et al., 2017; Rharass et al., 2014). Conversely, genetic disruption of respiratory chain components or mitochondrial quality-control factors (for example AIF and SURF1) in mouse NSCs or human neural progenitors impair proliferation and neuronal differentiation. Thus, ensuing, *in vivo* models recapitulate key features of related neurometabolic mitochondrial encephalopathies such as Leigh syndrome, including abnormal cortical cytoarchitecture (Inak et al., 2021; Khacho et al., 2016, 2017). These observations provide a conceptual framework in which *HPDL*-related disease might reflect not only a static developmental defect but also a dynamic failure of mitochondrial support for neurogenesis and long-term neuronal maintenance.

In this context, our previous study, investigating the consequences of *HPDL* loss-of-function in human neuroblastoma (SH-SY5Y) cells and HPDL Patient-derived cortical cultures (Baggiani et al., 2026), showed that HPDL is important for respiratory chain supercomplex (RCS) assembly, basal respiration, and redox balance. Indeed, mitochondrial ROS were elevated and CoQ₁₀ levels were paradoxically increased, suggesting possible mislocalization or inefficient utilization of this cofactor. In parallel, induced pluripotent stem-cell (iPSC) lines from four patients with *HPDL*-related spastic paraplegia (SPG83) were differentiated into 2D and 3D cortical *in vitro* models. These cultures exhibited premature neurogenesis of cortical progenitors at early stages (DIV16–30), with up-regulation of neurogenic gene programs and depletion of proliferative pools, leading to reduced organoid growth with a microcephaly-like appearance. Mitochondrial analyses in DIV16-30 progenitors revealed impaired RCS assembly, altered mitochondrial membrane potential, increased ROS, and dysregulated CoQ₁₀ content, directly linking *HPDL* deficiency to mitochondrial dysfunction in human neural cells. Importantly, pharmacological treatments with either 4-HB or mitochondrial-specific superoxide scavenger MitoTEMPO partially rescued the premature-neurogenesis phenotype in a strict mutation-dependent manner (Baggiani et al., 2026).

Together, these findings suggest that *HPDL*-related disease leads to an early disturbance of cortical neurogenesis driven by mitochondrial dysfunction. However, several key questions remain unanswered. First, it is unclear whether the disorder is purely neurodevelopmental or whether there is a progressive neurodegenerative component unfolding from late stages of cortical maturation onward. Second, the contribution of non-neuronal cell types—particularly astrocytes and oligodendrocytes, which are critical for synaptic support, glutamate and ion homeostasis, and metabolic coupling to neurons—has not been explored in *HPDL*-deficient human brain tissue (Molofsky and Deneen, 2015; Shi et al., 2025).

In the present study, we delved into HPDL biology, trying to address these gaps *via* long-term iPSC-derived human cortical cultures we had generated (Baggiani et al., 2024a, 2026). To do this, we combined quantitative immunofluorescence and bulk RNAseq at late differentiation stages (DIV90–120), particularly focusing on (*i*) the composition of cortical neuronal subtypes, (*ii*) the level of synaptic maturation, (*iii*) the abundance and cellular status of astrocytes and oligodendrocytes, all of these combined with (*iv*) occurrence of apoptotic cell death in the different populations. Building on our previous work, this long term, longitudinal analysis reveals that *HPDL* deficiency in human cortical mature cultures is associated with a striking triad of phenotypes: progressive neurodegeneration with synaptic loss, profound glial dysfunction, and unexpected late reactivation of early neurogenic transcriptional programs.

## Results

### Transcriptomics analysis in late stage HPDL cortical cultures suggest unbalanced expression in genes related to cellular adhesion, extracellular matrix synthesis, neuronal morphogenesis, and synaptic function

In our first report on HPDL mutant cortical cultures, we observed early dysregulation of neurogenesis associated with mitochondrial failure, leading to decreased organoid size and increased number of CTIP2^+^ cells in patient-derived compared to control cells. Here, we expand our investigation, studying HPDL-associated phenotypes combined with later cortical development and neuronal maturation. We analyzed the transcriptional scenario in a time-point analysis of HPDL mutant cells compared to control counterparts. To have an unsupervised and unbiased survey on possible phenotypes occurring in mature patient-derived cultures, we analyzed the transcriptional scenario of DIV120 HPDL mutant cells compared to control counterparts. In particular, together with the discrete time-point analysis, we followed longitudinal gene expression trends over time in each HPDL and control line, from earlier (DIV16 and DIV30, coming from our previous study) to later stages (DIV120). Dynamic temporal analysis found 4,532 and 6,970 DEGs in control and patient-derived cell cultures, respectively (**Fig. 1A**). Searching for patient-specific DEGs, we focused on genes that were not in common with controls (3585 genes; **Fig. 1B**), most of which were protein-coding (78.15%; **Fig. 1C**). GO analysis defined that most represented categories were related to mitochondrial assembly and function (yellow), nucleoside metabolism (green), CNS developmental processes (light blue) and ribosomal proteins (pink; **Fig. 1D**), mirroring previous findings on HPDL early cultures (Baggiani et al., 2026). Then, to capture static differences between HPDL mutant and control counterparts at later stages of cortical development in snapshot-based fashion, we focused attention on transcriptomic profiles at DIV120 time point, identifying 306 DEGs. GO and PANTHER enrichment analysis showed significant downregulation of categories associated with cell–cell adhesion and extracellular matrix organization (such as *Homophilic adhesion via plasma membrane adhesion molecules*, *Cell-cell adhesion via plasma membrane adhesion molecules*, *Extracellular matrix*, *Cadherin signaling pathway*, and others). In addition, KEGG pathway analysis stressed possible downregulation of synaptic function, with enrichment of *Neuroactive ligand-receptor interaction* and *Calcium signaling pathway* categories (**Fig. 1E**).

**Figure 1:**
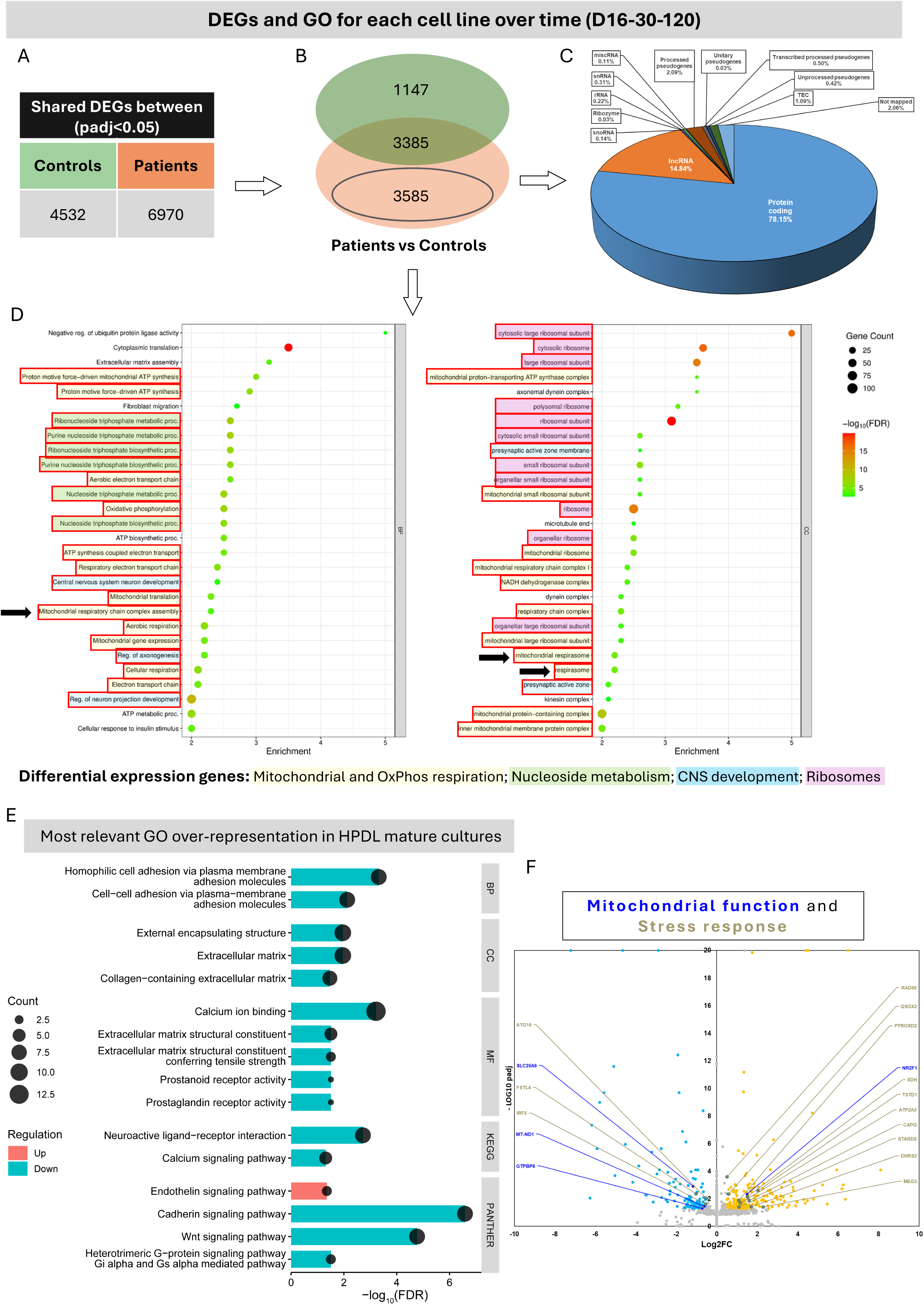
Longitudinal transcriptomic analysis reveals persistent molecular dysregulation in HPDL mutant cortical cultures. (A) Temporal profiling across DIV16, DIV30, and DIV120 identified 4,532 differentially expressed genes (DEGs) in controls and 6,970 DEGs in patient-derived cultures. (B, C) Among these, 3,585 DEGs were specific to patient-derived cultures and the majority was protein-coding transcripts (78.15%). (D) Gene Ontology enrichment analysis of patient-specific longitudinal DEGs highlighted pathways related to mitochondrial function and oxidative phosphorylation, nucleoside metabolism, CNS development, and ribosomal processes. (E) Analysis of DIV120 mature cultures showed predominant downregulation of gene categories associated with cell adhesion, extracellular matrix organization, and synaptic signaling. (F) Volcano plot highlighted the most dysregulated genes related to *Mitochondrial function* and *Stress response* groups in HPDL mature cultures.

To complement automated cluster analysis, we manually performed a literature-based gene-by-gene reassessment, allowing categorization of all DEGs via their known biological function in 8 main groups: *Neurodegeneration-associated genes*, *Astrocytic genes*, *Early developmental genes*, *ECM/ECM-associated effectors*, *Synaptic function*, *Neuronal morphogenesis/Axon guidance*, *Mitochondrial function*, and *Stress response* (summarized in Volcano plot; **Suppl. Fig. 1A**). Notably, downregulation of genes such *MT-ND1* and *SLC25A6*, coding for mitochondrial subunit of complex I and mitochondrial ADP/ATP antiporter, respectively, suggested mitochondrial dysfunction, similar to earlier stages (**Fig. 1F**). In the same way, increased expression of *QSOX2*, *DHRS2*, *XDH*, *STARD5*, and *PYROXD2* genes, combined with downregulation of *IRF5* gene, support persistence of oxidative stress in HPDL late cortical cultures (**Fig. 1F**).

Collectively, these findings support and corroborate our previous results linking disruption of cortical development and mitochondrial morpho-functional properties in HPDL patient-derived cortical tissues (Baggiani et al., 2026).

### Late stage HPDL cortical neurons show synaptic impairment

Based on abovementioned transcriptomic results, we performed immunostaining experiments to explore morpho-functional neuronal characteristics in HPDL mutant neurons at late stages.

To this end, we first evaluated SMI32 immunoreactivity (**Fig. 2A**), recognizing non-phosphorylated neurofilament heavy chain. This cytoskeletal marker is widely used to assess pyramidal neuron structural integrity and axonal health, and it is known to decrease under conditions of cellular stress and degeneration (BurianovÃ¡ et al., 2015; Hof et al., 1990; Ohm et al., 2025; Petzold, 2005; Voelker, 2004). The data showed an unchanged level of SMI32 mean fluorescence in HPDL derived late neurons compared to controls, except for Patient 3-derived neurons, in which a significant decrease was found (**Fig. 2B**). In addition, in order to characterize possible occurrence of neuronal maturation defects in HPDL mutant cultures, we evaluated the protein levels of RBFOX3 (a.k.a. NeuN) and MAP2, widely used as *bona fide* markers of neuronal maturation, *via* immunostaining in DIV90 patient-derived cortical tissue compared to control neurons. No changes were found in MAP2 mean fluorescence (**Suppl. Fig. 1B**), while the protein levels of transcription factor RBFOX3 showed a significant increase in Patient 2-derived neurons only (**Suppl. Fig. 1C**). Collectively, neuronal maturation in late HPDL mutant cultures seems to be comparable to control counterparts.

**Figure 2.**
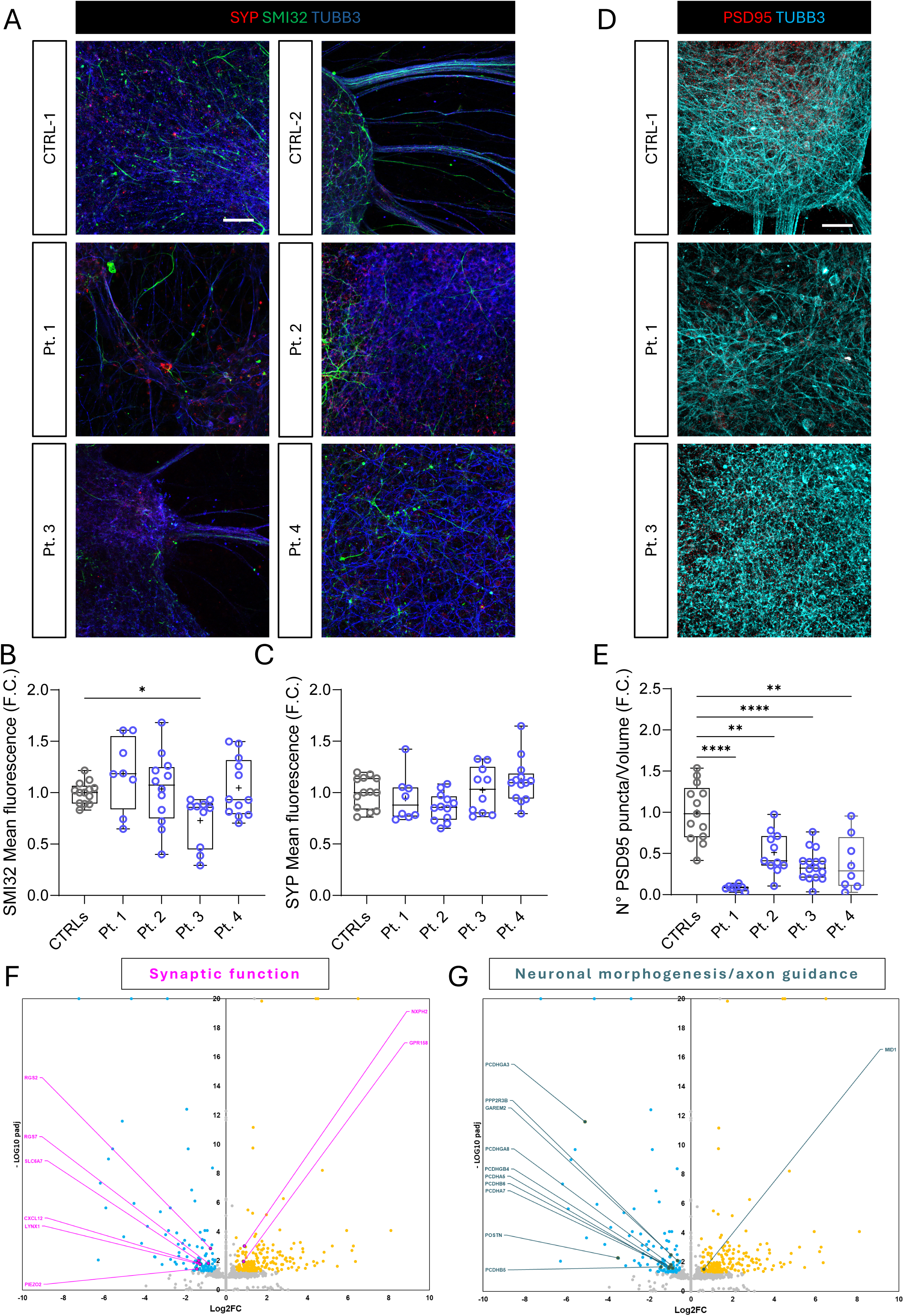
Late-stage HPDL mutant cortical neurons show impaired synaptic organization. (A) Representative confocal images of Synaptophysin (SYP), SMI32, and TUBB3 in DIV90 control and HPDL Patient 1 and 3 derived cortical neurons. (B, C) Box and whiskers blots showed the quantification of SMI32 and SYP mean fluorescence in control and HPDL mutant cultures. A significant reduction was found only for SMI32 mean fluorescence level in Patient 3-derived cultures. (D) Representative confocal images of PSD95 (post-synaptic marker) and TUBB3 in DIV90 control and HPDL Patient 1 and 3 derived cortical neurons. Scale bar: 50 µm. (E) Box and whiskers plot showed the quantification of PSD95 puncta density revealed a drastic reduction of excitatory post-synaptic puncta in HPDL neurons. (F, G) Volcano plots showing dysregulated genes associated with synaptic function and neuronal morphogenesis/axon guidance, supporting impaired synaptic and morpho-structural integrity in HPDL mutant neurons. (B, E) Welch’s ANOVA test, *post-hoc* Dunnett’s T3 multiple comparisons test; * *p*-value < 0.05, **** *p*-value < 0.0001. (C) One-way ANOVA test, *post-hoc* Holm-Šídák’s multiple comparisons test; p-value > 0.05. (B, C, E) All data are showed in Box and whiskers plots, expressed as fold change (F.C.) normalized to untreated controls, and represented as Min-to-Max. Mean is indicated with “+” symbol (N = 3 for each line). Scale bar: 50 µm.

Next, we focused on maintenance of the neuronal synaptic compartment. We therefore analyzed in DIV90 HPDL mutant and control neurons quantity and distribution of Synaptophysin (SYP) and PSD95, two universally accepted pre- and post-synaptic markers. While quantification of SYP mean fluorescence signal showed on average comparable levels (**Fig. 2C**), we observed a striking decrease in the number of PSD95 puncta (per volume) in HPDL mutant neurons derived from all different lines (**Fig. 2D, E**). These results corroborated the transcriptomic data on DIV120 neurons, where several genes involved in synaptic function and post-synaptic density organization as *CXCL12*, *SLC6A7*, *RGS2*, *RGS7*, *LYNX1* (**Fig. 2F**) were downregulated. Moreover, DEGs related to neuronal morphogenesis and neurite outgrowth (as *GAREM2*) were mostly downregulated (**Fig. 2G**), suggesting occurrence of morpho-structural defects.

These results consistently suggest that patient-derived neurons, despite retaining comparable levels of maturation markers, show a striking impairment of the excitatory post-synaptic compartment.

### Cellular composition in late HPDL patient-derived cortical cultures define unbalanced cell-type specification

The natural phenotypic outcome previously found in early HPDL mutant cortical cultures as a consequence of premature neurogenesis was the increase of deeper layer cortical neuron production. Following this paradigm, we expected either (*i*) a reduction of later-born upper layer neuron production if the neurogenesis process was interrupted or (*ii*) an increased/unchanged production if this neurodevelopmental dynamic were just accelerated, similarly to findings described by others (Shabani et al., 2023). To clarify this dichotomy, we analyzed cortical cell type composition in late (DIV90) HPDL mutant and control cortical cultures *via* immunostaining for CTIP2 (layer 5), and SATB1/2 (layers 2-3) (**Fig. 3A**). The normalized quantity of CTIP2^+^ neurons showed a decrease in Patient 1 neurons and an increase in Patient 3 and 4 derived cells (**Fig. 3B**), mirroring the results obtained in early HPDL mutant cortical cultures (Baggiani et al., 2026). In spite of this, no changes in the quantity of SATB1/2^+^ neurons occurred in Patient 2 and 3-derived cultures, while Patients 1-and 4-derived mature cortical tissues showed an increased number of upper layer neurons compared to control counterparts (**Fig. 3C**). This pattern seems to cast doubts on the hypothesis of interrupted neurogenesis, increasing the odds of alternative mechanisms such as accelerated corticogenesis.

**Figure 3.**
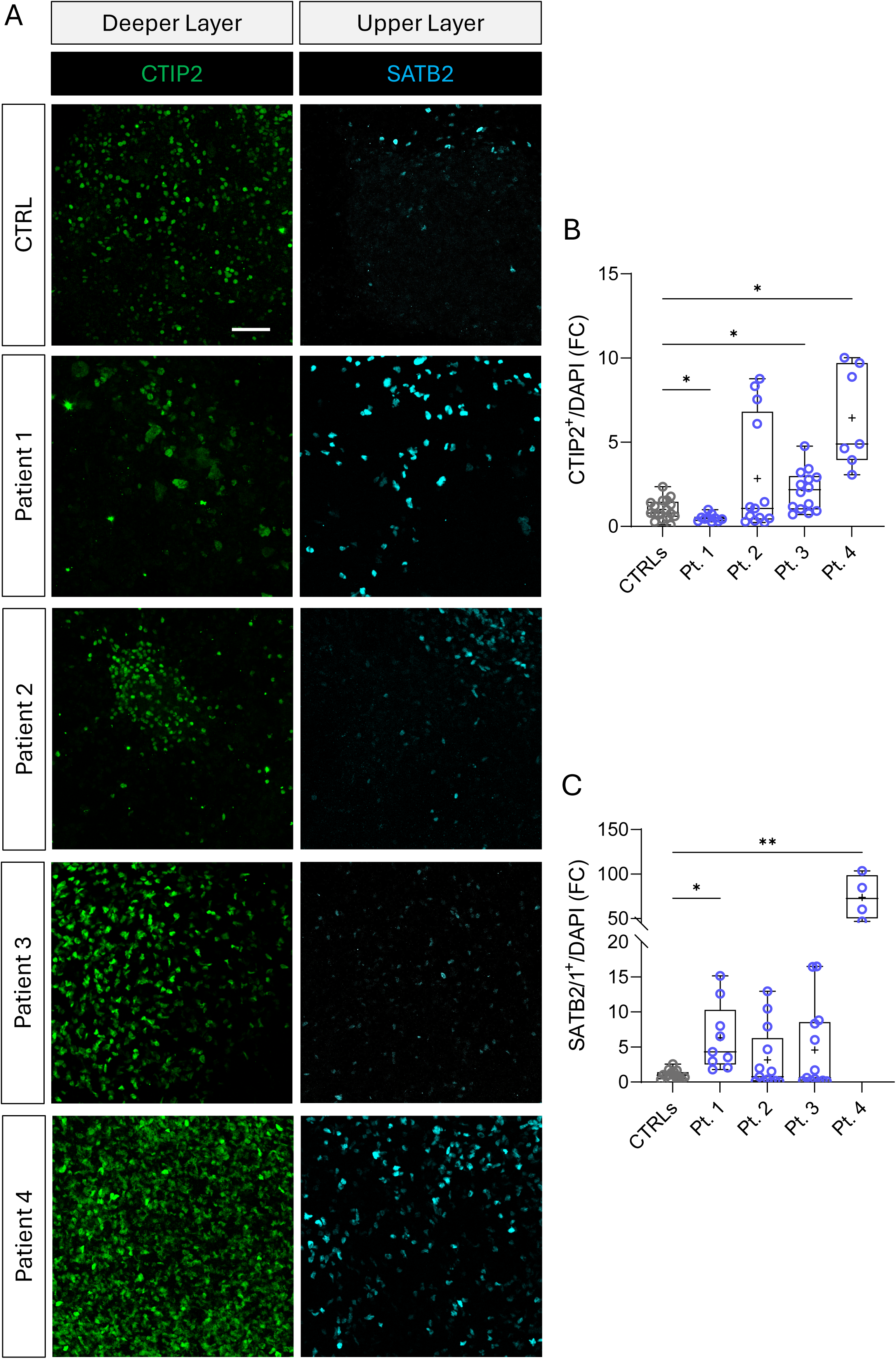
HPDL mutant cortical cultures display altered cortical layer composition at late differentiation stages. (A) Representative confocal images of DIV90 control and HPDL Patient-derived cortical cultures stained for CTIP2 (deeper-layer neurons), and SATB2/1 (upper-layer neurons). (B) Quantification of CTIP2^+^ cells showed patient-specific dysregulation of deeper-layer neuronal populations, with reduced levels in Patient 1 and increased levels in Patient 3- and Patient 4-derived cultures. (C) Quantification of SATB1/2^+^ cells showed increased upper-layer neuronal populations in Patient 1-and Patient 4-derived cultures, supporting an imbalance in cortical cell-type specification. (B) Welch’s ANOVA test, *post-hoc* Dunnett’s T3 multiple comparisons test; **** *p*-value < 0.0001. (C) Kruskal-Wallis test, *post-hoc* Dunn’s multiple comparisons test; *** *p*-value < 0.001. (B, C) All data are showed in Box and whiskers plots, expressed as fold change (F.C.) normalized to untreated controls, and represented as Min-to-Max. Mean is indicated with “+” symbol (N = 3 for each line). Scale bar: 50 µm.

In late cortical cultures, neural progenitors are known to switch from generating neurons to produce glia (Allen and Lyons, 2018). In this perspective, to complete our analysis on HPDL mutant late cortical populations we performed immunocytochemical staining in DIV90 patient-derived and control cortical cultures for glial populations, astrocytes (marked by GFAP) and oligodendrocytes (labelled by OLIG2; **Fig. 4A**). Surprisingly, the relative number of astrocytes was markedly reduced in patient-derived cultures, especially in Patient 1, in which this cell population was almost completely absent (**Fig. 4B**). Despite this, no signs of glial reactivity were observed; the few astrocytes present in HPDL cultures displayed a decrease, rather than an increase, in mean GFAP fluorescence (**Fig. 4C**). Although less evident, the same phenotype was also present for oligodendrocytes, as OLIG2^+^ cells were significantly reduced only in Patient 1 and 3 derived neurons (**Fig. 4D**). Notably, astrocytic impairment was also confirmed by transcriptomic analyses, in which astrocyte-specific genes such as *GJB6*, *P2RX5*, and *SLC22A3* were downregulated, while other more functional genes such as *SLC13A3*, *LPAR3*, *SLC5A12*, and *SLC26A9* were upregulated (**Fig. 4E**). In addition, consistent with categories enriched in GO and PANTHER analysis (**Fig. 1E**), extracellular matrix associated genes, usually expressed by astrocytes, were highly downregulated (**Fig. 4F**).

**Figure 4.**
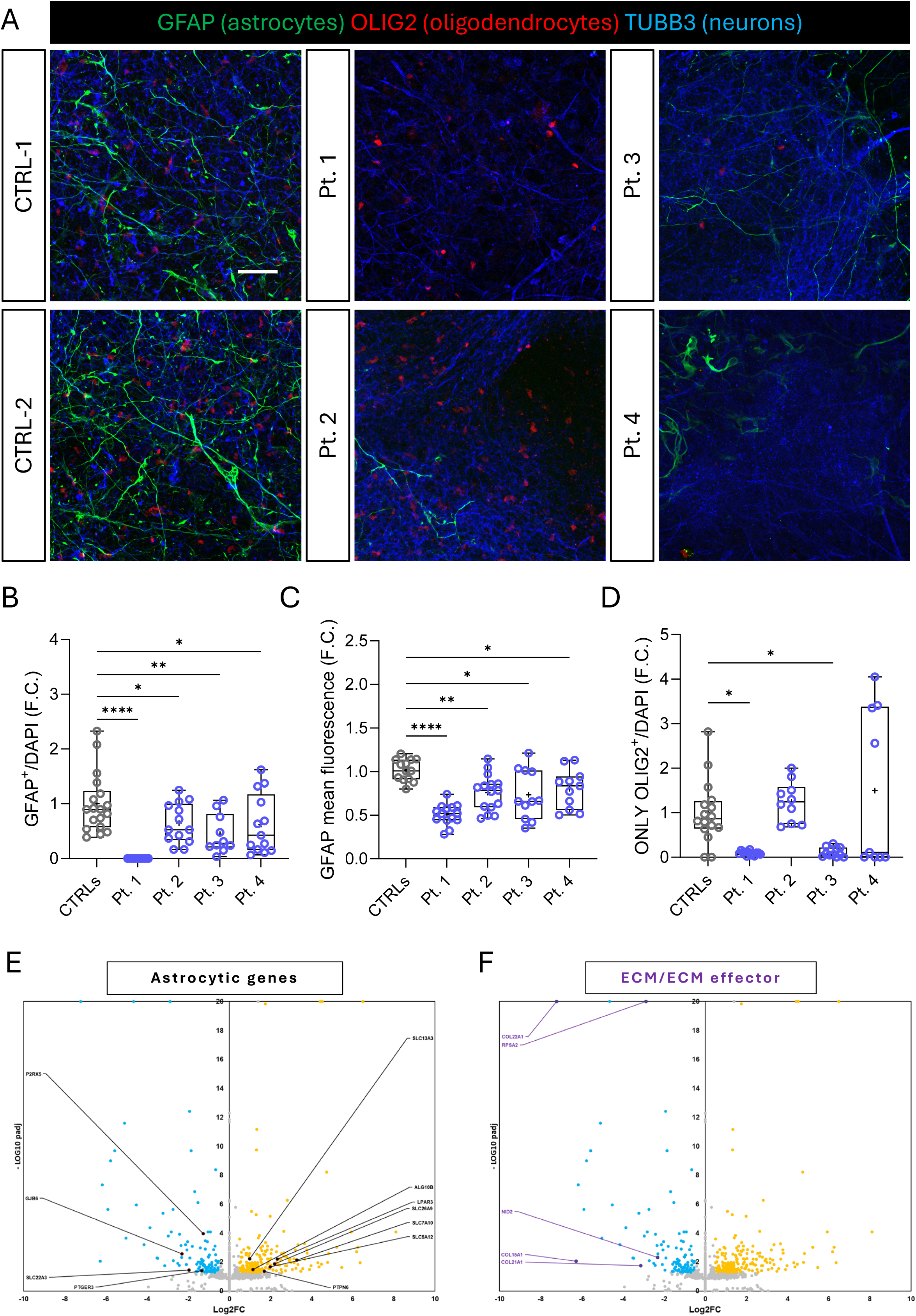
HPDL mutant cortical cultures show glial impairment and extracellular matrix dysregulation. (A) Representative confocal images of DIV90 control and patient-derived cortical cultures stained for GFAP (astrocytes), OLIG2 (oligodendrocytes), and TUBB3 (neurons). (B, C) Quantification of GFAP positive cells and mean fluorescence revealed a marked reduction of the astrocytic compartment in patient-derived cultures, particularly in Patient 1. (D) Quantification of oligodendrocyte (OLIG2^+^/GFAP^-^) showed significant alterations in Patient 1- and Patient 3-derived cultures. (E, F) Volcano plots highlighted the dysregulated astrocytic genes and ECM/ECM-associated effectors within the two related groups, supporting glial dysfunction and extracellular matrix impairment in mature HPDL mutant cortical cultures. (B) One-way ANOVA test, *post-hoc* Holm-Šídák’s multiple comparisons test; **** *p*-value < 0.0001. (C) Welch’s ANOVA test, *post-hoc* Dunnett’s T3 multiple comparisons test; **** *p*-value < 0.0001. (D) Kruskal-Wallis test, *post-hoc* Dunn’s multiple comparisons test; *** *p*-value < 0.001. (B, C, D) All data are showed in Box and whiskers plots, expressed as fold change (F.C.) normalized to untreated controls, and represented as Min-to-Max. Mean is indicated with “+” symbol (N = 3 for each line). Scale bar: 50 µm.

### Late HPDL mutant cortical cultures undergo neurodegenerative processes

A possible factor contributing to the phenotypic scenario ongoing in HPDL mutant cortical tissue at later stages of development and potentially explaining the observed imbalance in cell type composition, could be constituted by cell death. To this end, we stained DIV90 cortical cultures differentiated from patient-derived and control iPS lines, comparing the relative quantity of apoptotic cells (cCASP3^+^) within the astrocytic (GFAP^+^) and neuronal (TUBB3^+^) population (**Fig. 5A**). The data showed a significant enhancement of apoptosis in all HPDL Patient-derived cortical cultures, except for the ones deriving from Patient 2 (**Fig. 5B**). Interestingly, the rate of apoptotic astrocytes (cCASP3^+^GFAP^+^; **Fig. 5C**) in HPDL mutant cells did not differ from control counterparts (Patient 1 was excluded from the analysis since almost completely devoid of GFAP+ cells, as previously shown). Conversely, the percentage of apoptotic neurons (cCASP3^+^TUBB3^+^; **Fig. 5D**) were significantly increased in Patients 1, 2, and 3, while apoptotic cells negative for TUBB3 and GFAP markers (cCASP3^+^GFAP^-^TUBB3^-^; **Fig. 5E**) were found to be increased in Patients 1, 2, and 4. Consistently, a subset of genes form RNA-Seq analysis, such as *AATK*, *TRPV3*, *IRF6*, *GALNT9*, *FCGBP*, and *CNN2*, underwent dysregulation in DIV120 HPDL cultures (*Neurodegeneration-associated genes* cluster, **Fig. 5F**).

**Figure 5.**
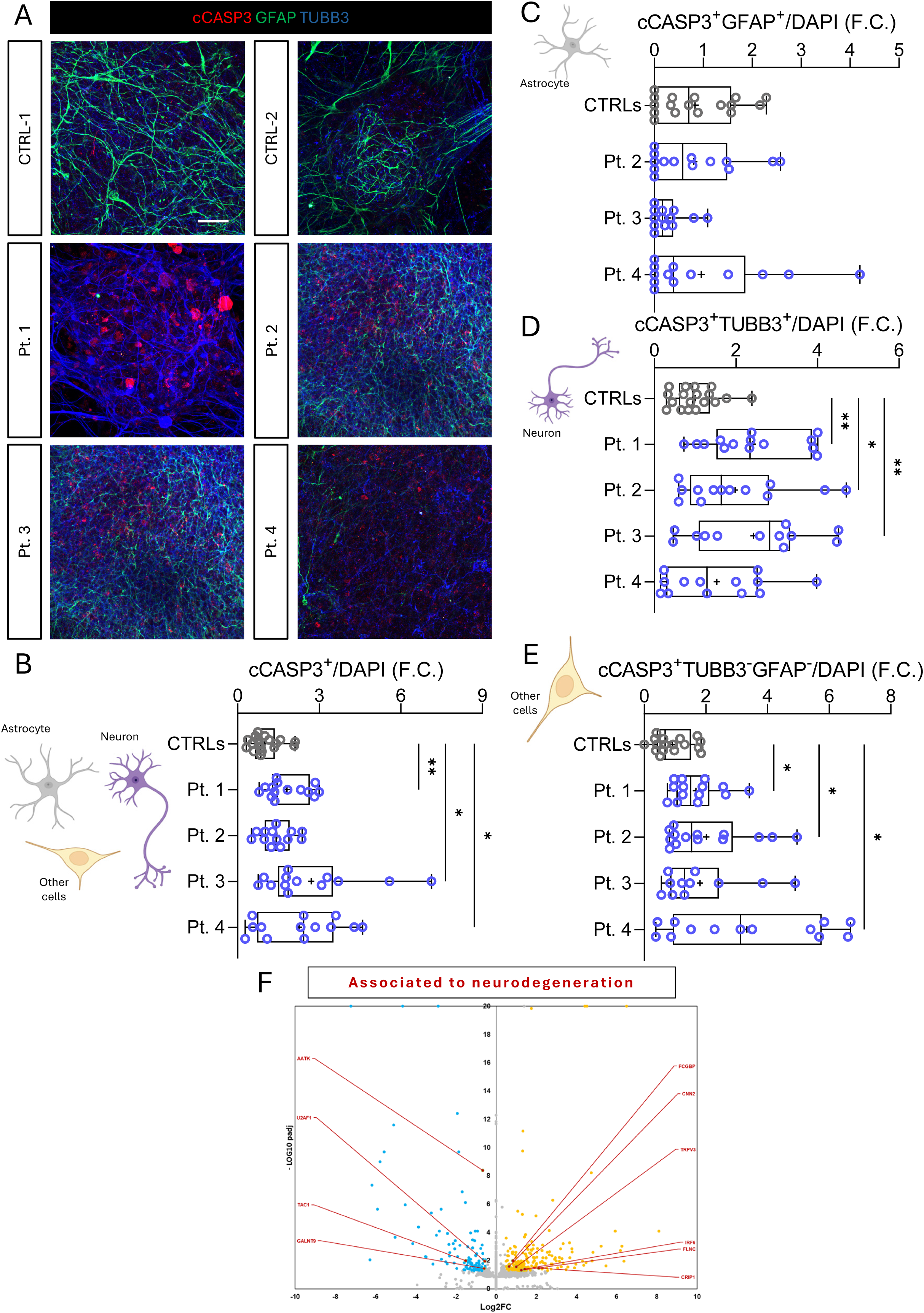
HPDL mutant cortical cultures undergo neurodegeneration at late differentiation stages. (A) Representative confocal images of cCASP3 (apoptotic cells), GFAP (astrocytes), and TUBB3 (neurons) in DIV90 control and HPDL cortical cultures. (B) Box and whiskers blot represented cCASP3^+^ cell quantification, showing an increased apoptosis in patient-derived cultures, with the exception of Patient 2. (C - E) Specific quantification of lineage-specific apoptotic cells (cCASP3^+^) among astrocytes (GFAP^+^), neurons (TUBB3^+^), and other cells (GFAP^-^/TUBB3^-^) showed an increased cCASP3^+^ neurons in Patients 1, 2, and 3, and increased cCASP3^+^ other cells in Patients 1, 2, and 4 derived late cortical cultures. (F) Volcano plot showed a dysregulated neurodegeneration-associated genes, including *AATK*, *TRPV3*, *IRF6*, *GALNT9*, *FCGBP*, and *CNN2*, in DIV120 HPDL mutant cultures. (B, E) Welch’s ANOVA test, *post-hoc* Dunnett’s T3 multiple comparisons test; ** *p*-value < 0.01, ** *p*-value < 0.01. (C) Kruskal-Wallis test, *post-hoc* Dunn’s multiple comparisons test; *p*-value > 0.05. (D) One-way ANOVA test, *post-hoc* Holm-Šídák’s multiple comparisons test; ** *p*-value < 0.01. (B - E) All data are showed in Box and whiskers plots, expressed as fold change (F.C.) normalized to untreated controls, and represented as Min-to-Max. Mean is indicated with “+” symbol (N = 3 for each line). Scale bar: 50 µm.

Collectively, these results suggest that reduction of astrocytic population in HPDL mutant cortical cultures is likely due to impairment of developmental mechanisms involved in astrogliogenic switch rather than triggered by cell death. Patient-derived cultures, in any case, exhibit a degenerative phenotype that seems to overlap with developmental processes.

### NEUROD4 ectopic expression in HPDL patient-derived mature cultures

Based on findings coming from our previous results (Baggiani et al., 2026) and present findings, all the data coming from transcriptomic profiling has shown highly predictive power in HPDL patient-derived *in vitro* model of corticogenesis. The last group of DEGs emerging from manual categorization of data from DIV120 RNA-Seq analysis surprisingly consists in *Early developmental genes*, comprising genes that are typically expressed during early cortical development. Notably, a subset of genes including *NEUROD4*, *NHLH2*, *LIN28B*, *LBX2*, *SOX3*, *PAX3*, and *PLAG1* displayed a markedly strong and heterochronic upregulation in late HPDL mutant cultures (**Fig. 6A**). This unexpected late-stage expression of early developmental genes suggested the possible occurrence of two putative scenarios: a) persistence of expression of early, pro-neural genes, counteracting astrogliogenesis switch; b) reactivation of neurogenic competence in astrocytes (also described as astrocyte reprogramming), known *in vivo* to be actually dependent on forced genetic expression of the NEUROD4 transcription factor (Masserdotti et al., 2015). To verify this last hypothesis, we performed a triple NEUROD4/GFAP/TUBB3 immunostaining on DIV120 mutant and control cortical cultures (**Fig. 6B**). Consistently with transcriptomics, a non-negligible proportion of astrocytes and neurons showed NEUROD4 positivity in cultures derived from all patients except Patient 2, whereas positive cells were highly sporadic and nearly absent in control counterparts. Despite its well-known function as a transcription factor with nuclear localization, NEUROD4 staining in our models was almost always cytoplasmic rather than nuclear, clearly demonstrating occurrence of a non-physiological condition (**Fig. 6C**). Notwithstanding, we quantified the proportion and identity of NEUROD4-stained cells. Strikingly, we found a strong increase (from four- to seven-fold) in astrocyte positivity, and an even sharper rise (at least five-fold) in the ratio of triple NEUROD4/GFAP/TUBB3 positive cells in DIV120 Patient 1, 3, and 4 derived cultures (**Fig. 6D**), actually endorsing the possibility of ongoing neuronal conversion by NEUROD4-expressing astrocytes. To further support our hypothesis, we quantified the number of OLIG2/GFAP double positive cells among the different conditions, since OLIG2 is strongly upregulated in cortical astrocytes only to block the induced reprogramming in neurons (**Suppl. Fig. 1D**) (Lai et al., 2026). Notably, an increase of OLIG2/GFAP positivity was only found in cultures deriving from Patient 2, consistently with absence of GFAP positivity in NEUROD4 population, and a reduction in Patient 1, probably related to astrocyte depletion (**Fig. 6E**).

**Figure 6.**
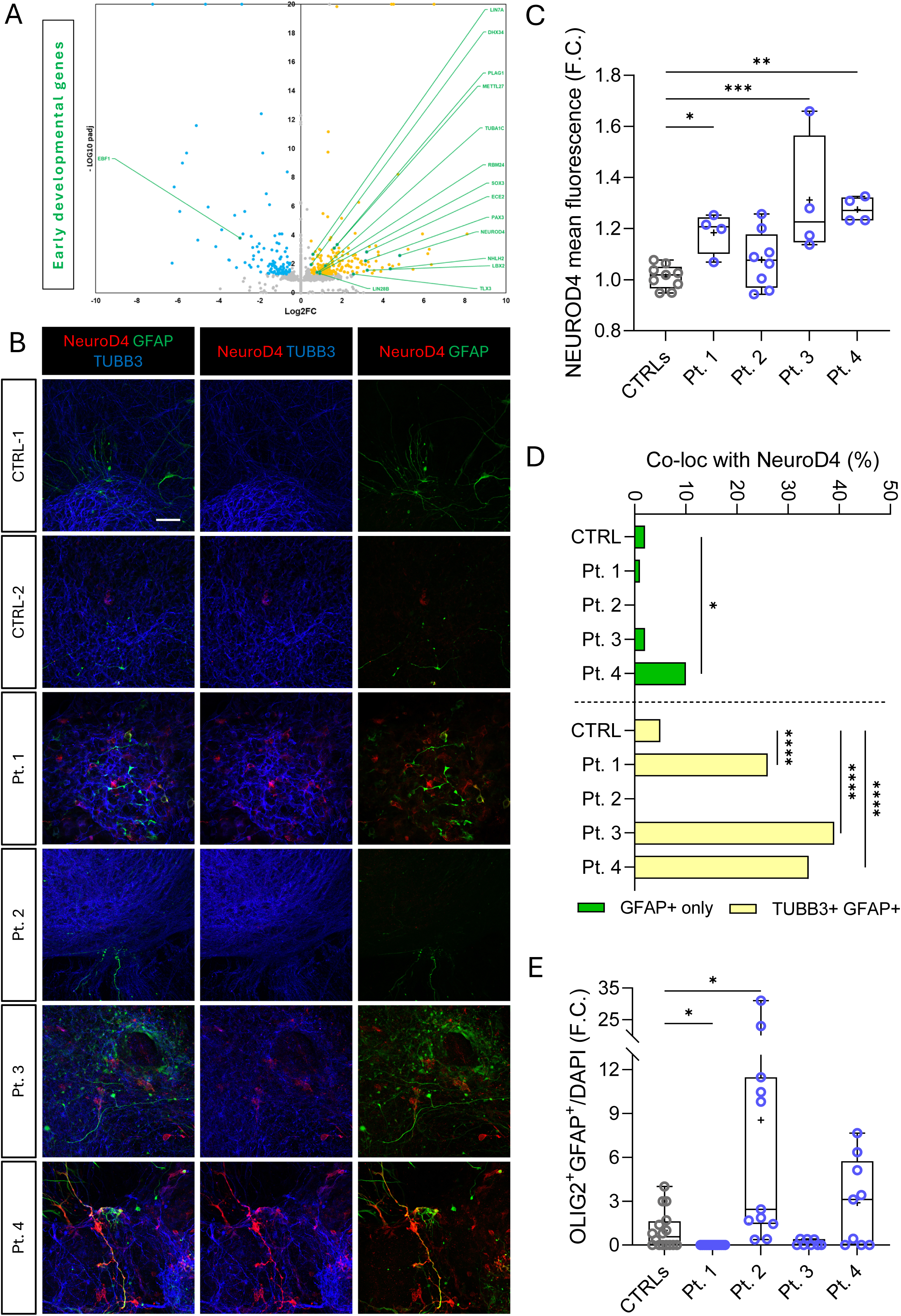
HPDL mutant mature cortical cultures show ectopic activation of early developmental genes. (A) Volcano plot showing dysregulation of early developmental genes, including *NEUROD4*, *NHLH2*, *LIN28B*, and *PLAG1*, in DIV120 HPDL mutant cultures. (B) Representative confocal images stained for NEUROD4, GFAP, and TUBB3 in DIV90 control and HPDL cortical neurons. (C) Box and whiskers plot showed the quantification of NEUROD4 mean fluorescence and the increased NEUROD4 signal in HPDL patient-derived cultures compared with controls, except Patient 2. (D) Co-localization analysis in NEUROD4^+^ population was represented in the bar plot, showing the increase of GFAP^+^ cells in Patient 4 and the steep rise of double GFAP and TUBB3 positivity in HPDL patient-derived cultures compared with controls, expect Patient 2. (E) Quantification of OLIG2/GFAP double-positive cells showed patient-specific alterations, with increased OLIG2/GFAP positivity in Patient 2-derived cultures and decrease in Patient 1. (C) One-way ANOVA test, *post-hoc* Holm-Šídák’s multiple comparisons test; *** *p*-value < 0.001. (D) two-tailed Z-test, comparing double NEUROD4/GFAP or triple NEUROD4/GFAP/TUBB3 positivity in CTRLs *vs* each HPDL Patient cultures. * *p*-value < 0.05, **** *p*-value < 0.0001. (C, E) All data are showed in Box and whiskers blots, expressed as fold change (F.C.) normalized to untreated controls, and represented as Min-to-Max. Mean is indicated with “+” symbol (N = 3 for each line). (D) All data are shown in histogram blot as rate of NEUROD4 population. (N = 3 for each line). Scale bar: 50 µm.

In conclusion, the expression of early cortical development genes, such as *NEUROD4*, in HPDL patient-derived mature cultures suggest new avenues in phenotypic interpretation of HPDL-related and other neurodevelopmental and neurodegenerative diseases.

## Discussion

In this work we extended our previous characterization of *HPDL* loss-of-function from early neural progenitor stages to late-maturing human cortical tissue, revealing that *HPDL*-related disease encompasses both neurodevelopmental and neurodegenerative components. Our earlier study established that *HPDL* deficiency impaired RCS assembly, elevated mitochondrial ROS, and drove premature neurogenesis with depletion of proliferative pools and reduced organoid growth (Baggiani et al., 2026). Here, using long-term iPSC-derived cortical cultures (DIV90–120) from the same Patients, we demonstrate that prematurely generated neurons subsequently undergo synaptic failure and apoptosis, in the context of severe astrocyte and oligodendrocyte loss and a re-activation of early neurogenic transcriptional programs. These findings suggest a model in which mitochondrial dysfunction induced by *HPDL* deficiency first distorts the tempo and balance of cortical neurogenesis and later compromises the maintenance of the neuronal and glial networks required for cortical integrity.

A key observation in our DIV90 patient-derived neurons is the significant dysregulation of cortical subtype markers. Specifically, we observed an increased density of CTIP2^+^ (layer 5) neurons in the late cultures of Patients 3 and 4 and a decrease in Patient 1, while the SATB1/2^+^ (upper layer) rate increased in neurons derived from Patients 1 and 4. Premature neurogenesis usually decreases neuronal numbers, but our findings point to an unanticipated paradox in mutant cortical cultures, with both deeper- and upper-layer neurons increased in number. A faster neurodevelopment process may allow neural progenitors to differentiate earlier while keeping their commitment could represent a possible explanation. In particular, we hypothesized an "IPC amplification model", in which mitochondrial dysfunction and elevated chronic oxidative stress, a peculiar hallmark in HPDL patient-derived cortical cultures, promote an increase in asymmetric divisions of RGCs, leading to a transient amplification of the intermediate progenitor cells (IPCs) during mid-neurogenesis period. Notably, this mechanism would be consistent with increase in asymmetric division occurring in the Aif-knockout mouse developing cortices, where mitochondrial dysfunction due to complex I impairment is a known driver of RGC-to-IPC fate switching (Apostolova et al., 2006; Khacho et al., 2016).

A transient increase of the IPC pool, swiftly followed by premature neurogenesis instead of amplifying the pool itself, would generate a higher number of neurons at the beginning, without apparent neural stem cell extinction. This model would fit perfectly with the time course of cortical differentiation in HPDL mutant cultures and is further supported by the metabolic switch occurring in IPCs, known to increase mitochondrial mass and use more OxPhos metabolism during neurogenesis (Brunetti et al., 2021; Fame and Lehtinen, 2021; Iwata and Vanderhaeghen, 2021; Khacho et al., 2016; Knobloch and Jessberger, 2017; Valenti and Vacca, 2023; Wanet et al., 2015). Interestingly, one of the most upregulated genes at RNA-Seq at later stages is *NHLH2*, a marker identified in a subset of human cortical IPCs (Pebworth et al., 2021). It is tempting to speculate that HPDL-mediated respiratory chain stress similarly perturbs NSC and IPC behavior, destabilizing neuronal identity at early and later stages via intense chronic ROS and metabolic signaling, as also supported by gene-expression changes over time. Future work will be necessary to delineate the precise mitochondrial mechanisms ongoing in this scenario.

Several findings from our data suggest that degenerative processes start in HPDL mutant cultures before cortical development completion, exacerbating the pre-existing neuro-developmental defects. First, late stages of HPDL Patient-derived cultures show a pronounced increase in cCASP3^+^ neurons and other non-neuronal cells compared with control, indicating ongoing apoptosis in differentiated cortical mature tissue.

Second, genes typically upregulated during neuronal stress and degeneration, including *AATK* (Tomomura et al., 2001), *TAC1* (Zhu et al., 2023), *FCGBP* (Gómez-Garre et al., 2022), and *CNN2* (Chou et al., 2025), were found in our patient cultures, together with DNA-repair and oxidative-stress–related genes such as *RAD50* (Lovell and Markesbery, 2007), *QSOX2* (Yan et al., 2023), and *PYROXD2* (Van Bergen et al., 2022). The convergence of apoptotic markers, DNA-damage signaling, and neurodegeneration-associated transcripts is consistent with chronic cellular stress culminating in neuron loss, aligning *HPDL*-related disease with other mitochondrial encephalopathies, where persistent energy and redox imbalance eventually trigger neurodegeneration (Anitha et al., 2023; Chung et al., 2011; Khacho et al., 2017; Panchal and Tiwari, 2019; Schwartz et al., 2007).

Third, even if maturation neuronal markers were unchanged, the density of PSD95^+^ excitatory postsynaptic puncta is drastically reduced across all HPDL Patient-derived cultures. This phenotype, associated with downregulation of synaptic genes involved in neurotransmission and modulation, supports impairment of synaptogenesis and impairment of synaptic function in neurons carrying *HPDL* pathological variants. This pattern resembles synaptic vulnerability described in primary CoQ deficiencies and Aif-deficient models, where mitochondrial defects in dendrites and synaptic compartments are sufficient to impair neurotransmission before neuronal death ensues (Leon et al., 2016; Millichap et al., 2024). Finally, it is known that mitochondria are also central to the regulation of synaptic function and neuronal survival. Synapses are energy-intensive structures that require local Ca²⁺ buffering and ATP production (Xavier et al., 2016), thus even subtle defects in mitochondrial OxPhos or ROS handling at synaptic terminals can lead to impaired neurotransmission and eventual synapse elimination, as extensively shown in models of Parkinson’s and Alzheimer’s disease (Cheng et al., 2010; Henchcliffe and Beal, 2008; Markesbery, 1997; Winklhofer and Haass, 2010; Xavier et al., 2016). The combination of synaptic loss, up-regulated degeneration markers, and apoptosis points to a progressive neurodegenerative process together with an anticipated neurogenesis.

These *in vitro* observations dovetail with clinical and imaging findings in patients and *Hpdl*-null mice. Clinically, *HPDL*-related disease often presents with progressive spastic paraparesis and ataxia, suggesting ongoing corticospinal and cerebellar tract degeneration (Alecu et al., 2025; Ghosh et al., 2021). Our human cortical dataset provides a cellular correlation for such persistent deficits, highlighting long-term synaptic and neuronal vulnerability in the context of *HPDL* deficiency.

The present work offers an additional novel level of interpretation. A previously unrecognized feature of *HPDL*-deficient cortical tissue is the severe alteration of glial populations. Our results showed a dramatic reduction in GFAP^+^ astrocytes across patient lines, particularly in Patient 1, where astrocytes are almost completely absent. Nevertheless, the few remaining cells display reduced GFAP mean fluorescence, suggesting the presence of cellular damage and/or limited capacity of a full cellular maturation. This phenotype does not align with many neurodegenerative disorders, where typical neurotoxic reactive astrocytes accumulate and actively contribute to neuronal death (Liddelow et al., 2017). Instead, our data point toward a failure of astrocytic development, likely compromising key homeostatic functions of the cortical tissue including extracellular-matrix (ECM) organization and structural support (Molofsky and Deneen, 2015; Rose et al., 2020), regulation of synaptogenesis (Dienel, 2013; Molofsky and Deneen, 2015; Sloan et al., 2017), glutamate uptake (Azarias et al., 2011; Rose et al., 2020; Zhang et al., 2023), ion buffering (Bélanger et al., 2011a; Chen et al., 2023), astrocyte-neuron lactate shuttle (ANLS) (Chen et al., 2023), and trophic support to neurons (Liddelow et al., 2017; Sloan et al., 2017).

On the other hand, oligodendrocyte-lineage impairment may also contribute to disease progression. Reduction of OLIG2^+^ cells in Patients 1 and 3 suggests a deficit in oligodendrocyte progenitors, which could impair myelination and axonal support, particularly in long corticospinal tracts that are characteristically affected in SPG83 (Damiani et al., 2024; Husain et al., 2020). In other HSP subtypes, mutations in mitochondrial or ER-shaping proteins (for example *SPG7* and *REEP1*; Damiani et al., 2024) lead to combined axonal and glial pathology, including crystalloid oligodendrogliopathy and abnormal myelin (Fünfschilling et al., 2012; Inak et al., 2021; Philips et al., 2021). Likewise, our results raise the possibility that *HPDL* deficiency disrupts glial lineages and that glial dysfunction is not merely secondary but may actively shape the neurodegenerative trajectory.

Occurrence of depletion in glial population is also supported by transcriptomic profiling, since genes typically associated with astrocytes and ECM organization are markedly dysregulated in patient cultures. The enrichment of GO categories related to ECM organization, cell–cell adhesion, prostaglandin, and neuroactive ligand-receptor signaling further indicates a possible remodeling of the perineuronal environment and glial–neuronal communication (Bélanger et al., 2011a). Given that astrocytes critically shape synaptogenesis and synaptic pruning respectively through both secreted ECM molecules and direct phagocytosis of synapses (Sloan et al., 2017), alterations in astrocyte number and ECM composition could likely be the main driver or at least vastly exacerbate the observed synaptic deficits, corroborated by RNA-seq findings. In this view, the observed maintenance of synaptophysin levels in pre-synaptic terminal only apparently contrasts with the decrease of PSD95 puncta in patient-derived cultures, as the rise of this marker is correlated with maturation of post-synaptic terminals (Essayan-Perez and Südhof, 2023; Shi et al., 2012).

Together with the direct effect due to strong downregulation of ECM/ECM effector genes, we can also speculate that ECM stability and morphology could be affected by increased acidification due to an excess of lactate release from cortical cells. In particular, neurons require a high energetic demand for their activity, thus they preferentially perform OxPhos respiration and Pentose Phosphate Pathway (PPP) (Cohen, 1950; Herrero-Mendez et al., 2009; Jimenez-Blasco et al., 2024), outsourcing glycolysis-mediated pyruvate production and lactate conversion to glial cells (Bolaños and Magistretti, 2025; Chen et al., 2023). In HPDL cortical tissue, impaired respiration likely leads mutant neurons to switch their energetic metabolism from OxPhos toward glycolysis and, consequently, lactate conversion for NAD^+^ regeneration (Chen et al., 2023; Mason, 2017). Several clinical reports showed a mild elevation of plasma lactate and pyruvate concentrations in patients harboring biallelic *HPDL* variants (Husain et al., 2020; Numata-Uematsu et al., 2021; Zeviani et al., 1996). Transcriptomics seems again to support this hypothesis, as the increase in energy demand could determine the observed upregulation of *SLC7A10*, coding for an aminoacid transporter expressed in neurons and astrocytes. Upregulation of *SLC5A12*, coding for an astrocytic low-affinity lactate importer for lactate, could instead reflect an attempt of astrocytes to buffer an excess of lactate from the ECM, as previously reported (Mächler et al., 2016).

In addition, while glial cells favor anaerobic metabolism, an increase in OxPhos activity can occur in some physiological or pathological conditions such as high energy demand (Hertz et al., 2007), higher neuronal activity (Dienel, 2013; González-Gutiérrez et al., 2020), hypoglycemia (Choi et al., 2012), lipid beta-oxidation (Mi et al., 2023), and metabolic stress (Zhang et al., 2023), making mitochondrial failure occurring in HPDL cortices a possible important modifier for these processes. A crucial physiological function of the astrocytic population consists of supplying antioxidant precursors to neurons (Bélanger et al., 2011a; Dringen et al., 1999). Imperfect mitochondrial supercomplex assembly naturally occurring in astrocytes actually leads to increased ROS formation, requiring an even more powerful ROS-scavenging machinery (Bélanger et al., 2011b; Cai et al., 2023; Jimenez-Blasco et al., 2015; Lopez-Fabuel et al., 2016). This phenomenon represents a part of a complex system necessary for neurons to scavenge the mitochondrial ROS generated by OxPhos respiration and Ca²⁺ buffering deriving from synaptic activity (see review of Bolaños and Magistretti, 2025 for detailed information). Notably, mitochondrial dysfunction has been implicated in several neurodegenerative and neurodevelopmental disorders, sometimes even preceding neuronal pathology (Mi et al., 2023).

Taken together, these findings could redefine *HPDL*-related disease as a disorder of the entire cortical neuro–glial unit. Premature impaired neurons presumably enter a hostile microenvironment characterized by loss of astrocytic and oligodendrocytic support, ECM disarray, and chronic stress signaling, hampering synaptic maturation and easing apoptotic cell death. Future studies using cell-type-specific manipulations or co-culture systems will be required to dissect the relative contributions of intrinsic neuronal vulnerability versus glial defects.

Finally, one of the most unexpected observations in our late-stage cortical cultures is the robust expression of early developmental transcription factors and progenitor-associated genes. Bulk RNA-seq at DIV120 reveals actual upregulation of genes as *SOX3*, *LIN28B*, *NEUROD4*, *LBX2*, and other early neurogenic regulators in patient-derived neurons compared with controls. The late activation of early neurogenic programs underscores that *HPDL* deficiency perturbs not only the timing of the initial neurogenic wave (premature neurogenesis at DIV16; Baggiani et al., 2026), but also the stability of cell identity at much later stages. Notably, LIN28B, NEUROD4, and PLAG1 transcription factors are all reported to bias toward neurogenic over gliogenic fate (Balzer et al., 2010; Oh et al., 2017). Overexpression of Plag1 in the mouse developing cortex, in particular, has been shown to promote neuronal differentiation at the expense of astrocyte production (Gasperoni et al., 2024; Sakai et al., 2019), an effect that could fit well with the observed phenotypes of prolonged neurogenesis and impaired gliogenesis. These transcriptional and protein-level changes suggest a late engagement of neurogenic programs that normally operate only at early progenitor stages.

A few, not mutually exclusive, interpretations can be considered. First, re-expression of early developmental genes has been shown in several cases to occur in mature neurons right before acquiring frank degenerating phenotypes (Andorfer et al., 2005; Folch et al., 2011; Frost, 2023; Namboori et al., 2021; Yang et al., 2006). Second, in other vertebrate systems, such as adult newt brain and neurogenic niches of the mammalian hippocampus, ROS-dependent activation of progenitor cells after injury triggers neurogenesis and partial functional recovery (Le Belle et al., 2011; Hameed et al., 2015; Nakatomi et al., 2002). Following this perspective, upregulation of neurogenic genes could reflect a compensatory attempt of resident cortical astrocytes to regenerate neurons in response to ongoing degeneration. This hypothesis would nevertheless be consistent with the observation that NEUROD4, a transcription factor normally required for initial neuronal differentiation (Hardwick and Philpott, 2015) and fundamental in astrocyte-neuron direct reprogramming (Masserdotti et al., 2015; Wang et al., 2023), was surprisingly found in astrocytes in DIV120 mutant cultures. In addition, a substantial subset of cells was also co-localized with neuronal markers, suggesting the occurrence of transitional neuro–glial states. In support of this hypothesis comes results in Patient 2, with positivity for OLIG2 inversely correlated with the absence of NEUROD4 expression. Besides confirming the observed diversity existing among cortical cultures harboring different *HPDL* pathological variants (Baggiani et al., 2026), these data would actually fit well with a model proposed in a recent report showing that expression of OLIG2 consistently rises in astrocytes as an “innate” barrier to counteract this cellular conversion (Lai et al., 2026). In any case, it should be noted that NEUROD4 had almost always a cytoplasmic rather than nuclear localization in *HPDL*-deficient cultures, maybe indicating the occurrence of abortive neurogenic response due to aberrant trafficking or sequestration. In addition, chronic stress could have a huge impact since the apparent reactivation of neurogenic programs occurs in a context of profound astrocyte loss, ECM remodeling, and mitochondrial dysfunction during neurogenesis (Iwata and Vanderhaeghen, 2021; Khacho et al., 2017; Liddelow et al., 2017; Molofsky and Deneen, 2015; Sloan et al., 2017), conditions that are unlikely to sustain effective neurogenesis.

The co-occurrence of mitochondrial gene dysregulation, oxidative-stress markers, and early developmental transcription factors in our dataset is consistent with this scenario.

Notably, a possible link between glial reprogramming to neurons mediated by NEUROD4 and glutathione-based antioxidant axis has been recently explored, posing interest for future progress in defining the possible effect of chronic oxidative stress in cellular behavior in cortical cells *in vivo* (Wang et al., 2023).

## Conclusion

The findings described in this work collectively redefine HPDL-related disease as a disorder of the entire cortical neuro–glial unit. Premature impaired neurons presumably enter a hostile microenvironment characterized by loss of astrocytic and oligodendrocytic support, ECM disarray, and chronic stress signaling, hampering synaptic maturation and easing apoptotic cell death. Studies in co-culture systems will be required to dissect the relative contributions of intrinsic neuronal vulnerability versus glial defects.

By integrating early progenitor-stage data with late-stage cortical phenotype, our study provides a temporal framework for *HPDL*-related cortical pathology. We propose that HPDL deficiency triggers a cascade in which (*i*) mitochondrial dysfunction and ROS promote premature neurogenesis and glial depletion, (*ii*) the resulting cortical tissue lacks sufficient glial support and exhibits intrinsic synaptic vulnerability, leading over time to (*iii*) synaptic loss, neuronal apoptosis, and re-activation of early neurogenic programs, as schematized in **Suppl. Fig. 1E**. This model reconciles the neurodevelopmental and neurodegenerative aspects of *HPDL*-related disease and opens several potential therapeutic windows: early interventions targeting mitochondrial function (for example 4-HB/4-HMA and antioxidants) and later strategies aimed at preserving synapses and glial function.

Obviously, our study has limitations. All data are derived from *in vitro* human cortical cultures, which cannot fully recapitulate *in vivo* circuit integration, vascularization, or systemic influences. The generation of cortical organoids from HPDL mutant lines could partially solve these issues. In addition, this study would also benefit from the analysis of an isogenic control line, although we clearly consider phenotypic diversity between HPDL mutant lines as a typical feature occurring in this disease.

Despite these caveats, the convergence of neurodegenerative, glial, and developmental signatures in HPDL patient-derived cortical tissue provides a coherent picture of a progressive cortical failure rooted in mitochondrial dysfunction. Future work combining single-cell multi-omics, live-cell imaging of mitochondria and synapses, and cell-type specific rescue experiments will be essential to test the proposed mechanisms and to evaluate whether CoQ head-group intermediates or targeted mitochondrial therapies can modify not only early developmental trajectories but also late neurodegenerative outcomes in *HPDL*-related disease.

## Materials and Methods

### Cell Reprogramming

Human dermal fibroblasts (HDFs), obtained from patient skin tissue via punch biopsies, were amplified in HDF medium and reprogrammed as previously described (Baggiani et al., 2024b; Okita et al., 2013; Schmitt et al., 2017). From about 14 days onwards, iPS “islands” appeared, and medium was switched to StemFlex medium (A3349401, Thermo Fisher Scientific). iPS clones with uniform flat and round shaped morphology were picked in sterile conditions and propagated as single cell lines. After seven passages, all iPS clones were characterized as described (Baggiani et al., 2024a, 2024b).

### Cell culture

iPSCs and neurons were cultured in standard conditions at 37 °C and 5% CO_2_. ACS1019 iPSCs were bought from ATCC, Manassas, VA, USA. Human iPSCs were cultured on Geltrex coated 6-well plates in StemFlex medium, refreshing medium every other day. Cells were passed every 5-7 days with ReLeSR™ (100-0484, Stem Cell Technologies), following manufacturer instructions. Cells have been differentiated following the dual-SMAD inhibition protocol (Chambers et al., 2009; Maroof et al., 2013). Briefly, pluripotent cells were harvested at high confluence on Geltrex and cultured for 12 days in neural induction medium containing 100 nM LDN193189, 10 µM SB431542, and 2 µM XAV939 (72147, 100-1051, 72674 respectively, Stem Cell Technologies), splitted with Accutase (A1110501, Thermo Fisher Scientific) and replated on poly-D-lysine/laminin coated coverslips (A3890401, Thermo Fisher Scientific and L2020, Sigma, respectively). From day 16 onward, medium was switched to terminal differentiation medium containing 30 ng/ml BDNF (AF-450-02, PeproTech) to improve neuronal differentiation and maturation. Differentiated cells were harvested at day *in vitro* (DIV) 90 and DIV120 and processed for immunostaining (see below) or RNA extraction via RNeasy Plus kit (74134, Qiagen).

### PSD95 puncta quantification

We counted the number of PSD95 puncta per volume unit (1 um^3^). To do this, images were thresholded keeping the values constant inside every differentiation, we performed the “watershed” function of Fiji and we counted the puncta with the Fiji plugin “analyze particles”, setting a range size (um^2^) of 0.3-Infinity and circularity from 0 to 1 for every line in every differentiation. Then, we normalized the ratio between the number of PSD95 puncta and volume (picture area *x* stack number) on the CTRL line of every differentiation.

### Cell counting method

Using Fiji plugin “cell counter”, we calculated the number of cells positive for CTIP2, SATB2/1, GFAP, OLIG2 (both positive or negative for GFAP), cCASP3 (total positivity, with or without TUBB3, and with or without GFAP), and NEUROD4 population. Then, we normalized the ratio between the counted cells and total cells (DAPI) on the CTRL line of every differentiation, except for NEUROD4, where the normalization was calculated on total number of NEUROD4 positive cell.

### Fluorescence quantification

Fiji software was used to analyze the mean fluorescence of SMI32, SYP, GFAP, NEUROD4, RBFOX3, or MAP2. In particular, images were thresholded keeping the values constant inside every differentiation and used to select ROIs. Then, the mean fluorescence of the selected marker was measured inside the ROI. Every measure was normalized on the CTRL line of every differentiation.

### Statistical analysis

All data in the manuscript represent three or more independent experiments. We performed the statistical analysis using GraphPad Prism 9.0.0 software. All data were analyzed using either parametric or non-parametric methods, based on the normal distribution analysis. For parametric data, we additionally tested whether standard deviations (homoscedasticity) were significantly different across groups. The significance among groups with 1 independent variable was measured with one-way ANOVA, Welch’s ANOVA, or Kruskal-Wallis tests. About NEUROD4 population, we used two-tailed Z-test, comparing double NEUROD4/GFAP or triple NEUROD4/GFAP/TUBB3 positivity in CTRLs *vs* each HPDL Patient cultures. Statistical significance is reported as: * P-value ≤ 0.05, ** P-value ≤ 0.01, *** P-value ≤ 0.001, or **** P-value ≤ 0.0001.

### Immunostaining, cell counting, and mean fluorescence analysis

Different cell types were treated in the same way, fixing with 4% formaldehyde (157-4-100, Electron Microscopy Science) for 12 min at RT and permeabilizing in PBS with 0.5% Triton X-100 (X100, Sigma) for 10 min at RT. PBS with 5% Normal Goat Serum (NGS; S-1000, Vector Laboratories) and 0.3% Triton X-100 was used as blocking solution for 1 h at RT and incubated with primary antibodies (SYP, 1:100, 36406, Cell signaling; TUBB3, 1:200, ab41489, Abcam; CTIP2, 1:500, ab18465, Abcam; SMI32, 1:300, ENZ-ABS219-0100, Enzo; SATB2/1, 1:400, ab51502, Abcam; cCASP3, 1:500, 9661, Cell Signaling; PSD95, 1:1000, ab18258, Abcam; OLIG2, 1:100, ab109186, Abcam; GFAP, 1:400, g3893, Sigma; NEUROD4, 1:500, pa5-115642, Thermofisher; RBFOX3, 1:500, ABN78, Millipore; MAP2, 1:200, ZMS1013, Sigma) in antibody solution (PBS with 3% NGS and 0.2% Triton X-100) at 4 °C overnight. Cells were then washed 3 times with PBS and incubated in the same antibody solution with secondary antibodies (Goat anti-rabbit IgG (H+L) 555, 1:500, a21429, Invitrogen; Goat anti-mouse IgG (H+L) 647, 1:500, a21236, Invitrogen; Goat anti-chicken IgG (H+L) 647, 1:500, a32933, Invitrogen; Goat anti-mouse IgG (H+L) 488, 1:500, a11029, Invitrogen; Goat anti-rat IgG (H+L) 488, 1:500, a11006, Invitrogen) for 1 h at RT. All nuclei were counterstained with DAPI (1 μg/ml; 10236276001, Merck). All images were acquired with a Zeiss LSM 900 confocal microscope (Zeiss) and processed with Fiji software (ImageJ 1.54f).

### Transcriptomics analyses

RNA quality control was performed on 48 RNA samples. RNA integrity was assessed using the RNA 6000 Nano Kit on a Bioanalyzer (Agilent Technologies). RNA samples were quantified using the Qubit RNA BR Assay Kit on a Qubit fluorometer (Thermo Fisher Scientific). All samples met the requirements for RNA sequencing. Illumina RNA-seq libraries were generated using the TruSeq stranded mRNA ligation kit (Illumina) from 400 ng of RNA samples, after poly(A) capture and according to manufacturer’s instructions. Quality and size of RNAseq libraries were assessed by capillary electrophoretic analysis with an Agilent 4150 Tape station (Agilent). Libraries were quantified by real-time PCR against a standard curve with the KAPA Library Quantification Kit (KapaBiosystems, Wilmington, MA, USA). Illumina Sequencing - Libraries were pooled at equimolar concentration and sequenced in 150PE on a NovaSeq6000 (Illumina) generating on average 29.8 million fragments per sample.

Quality of reads was assessed using FastQC software (http://www.bioinformatics.babraham.ac.uk/projects/fastqc/). Raw reads were trimmed with fastp (v.0.21.0) (Chen et al., 2018) with --trim_poly_x parameter, to remove adapters and low-quality bases with default parameters. Filtered reads were aligned to the Homo sapiens reference genome (Ensemble version 109) using STAR aligner (v2.7.9a) with parameter --peOverlapNbasesMin 5. Reads distribution on CDS, intronic and intergenic was computed using reads_distribution.py script from RSeQC suite (v4.0.0) (Wang et al., 2012). Gene expression quantification has been performed using RSEM and the Homo sapiens Ensembl v.111 annotation on the two different time-points and over-time. Genes-level abundance estimated counts and gene length obtained with RSEM (v1.3.1; https://github.com/deweylab/RSEM) were summarized into a matrix using the R package tximport (within DESeq2 package) and subsequently the differential expression analysis was performed with DESeq2 v.1.38.3 (Love et al., 2014). To generate more accurate log2 fold change estimates for low expressed genes, the shrinkage of the Log2 FoldChange was performed applying the apeglm method (Zhu et al., 2019). For transcriptomic analysis of patient-derived cortical neurons at DIV120, GO enrichment and gene clustering for up- and downregulated categories was performed via ShinyGO (Szklarczyk et al., 2023), keeping for both padj< 0.05 and LFC >|0.5|. Chord and bar plots were generated using SRplot (Tang et al., 2023).

## Acknowledgements

We thank all the members of Molecular Medicine Lab for overall theorical and practical scientific support.

## Supplementary material

Supplementary material is available online.

---

**Supplementary Figure 1.** Integrated transcriptomic, neuronal maturation, glial, and mechanistic overview of late HPDL mutant cortical pathology. (A) Volcano plot summarizing categorization of DIV120 differentially expressed genes in the different groups: *Neurodegeneration-associated genes*, *Astrocytic genes*, *Early developmental genes*, *ECM/ECM effectors*, *Mitochondrial function*, *Neuronal morphogenesis/Axon guidance*, *Stress response*, and *Synaptic function*. (B, C) Quantification of neuronal markers showed unchanged MAP2 mean fluorescence and a Patient 2-specific increase in RBFOX3 mean fluorescence, indicating overall preserved neuronal maturation marker expression in late HPDL mutant cultures. (D) Representative confocal images of OLIG2 and GFAP immunostaining, focusing on OLIG2/GFAP double positive cells in control and Patient 2-derived cultures. (E) Schematic model proposing that HPDL deficiency links mitochondrial impairment to glial loss, ECM disruption, synaptic vulnerability, cell death, and late reactivation of early neurogenic programs. (B, C) Welch’s ANOVA test, *post-hoc* Dunnett’s T3 multiple comparisons test; *p*-value > 0.05, ** *p*-value < 0.01. All data are showed in Box and whiskers plots, expressed as fold change (F.C.) normalized to untreated controls, and represented as Min-to-Max. Mean is indicated with “+” symbol (N = 3 for each line). Scale bar: 10 µm.

