## Supplementary figures and images for "Pathological variants in *HPDL* cause collapse of the neuro-glial unit during human cortical maturation"

### Supplementary Figure 1

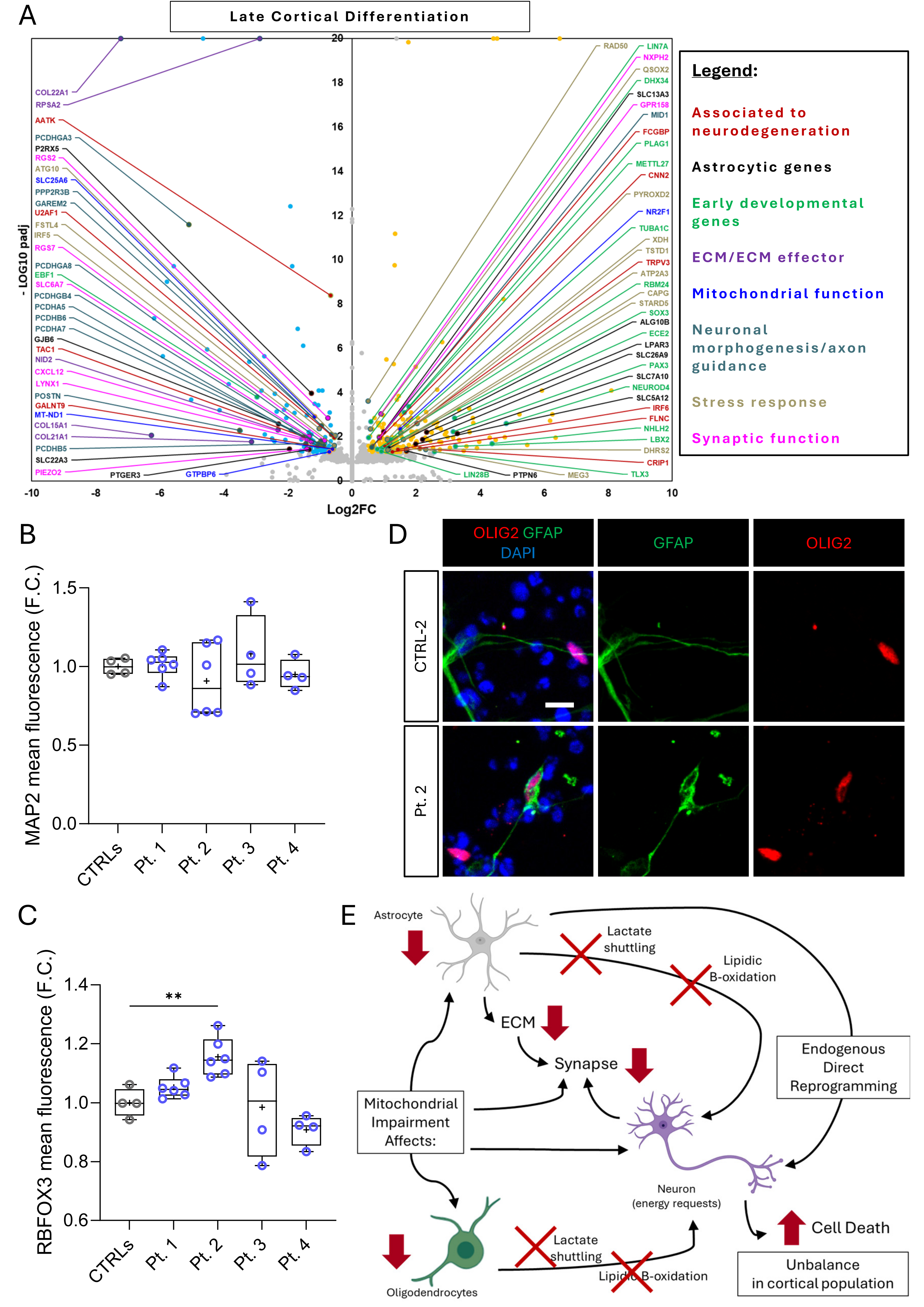
